# ButterflyVI: enabling high-throughput variant interpretation and biomarker discovery with functional genomics

**DOI:** 10.64898/2026.01.20.700339

**Authors:** Debora Sesia, András Hatos, Pasquelena De Nittis, Giovanni Ciriello

## Abstract

Characterizing the functional and therapeutic relevance of cancer mutations is a primary goal and a major challenge in precision oncology. Whereas predictive approaches exist, they often lack functional validation, which cannot scale to large numbers of cancer-associated variants. Here, we integrated statistical learning with experimental evidence from high-throughput functional screenings to annotate >20000 unique variants. We defined a variant as functional if it altered the effect of gene loss, and classified four possible outcomes: oncogene and tumor suppressor dependencies, mutation tolerance, and bypass-of-essentiality. Up-to-60% of variants annotated as functional were previously considered of unknown significance. Paradoxically, bypass-of-essentiality was common among loss-of-function (LoF) variants at several tumor suppressors, including *VHL*, *ARID1A*, and *RBM10*. In these cases, loss of the wild-type genes was deleterious, independently of the tissue of origin, but not when they already harbored recurrent LoF variants, suggesting loss of these tumor suppressors provides a context-specific advantage. Using our annotations, we discovered somatic variants that increased sensitivity to loss of therapeutically targetable genes, representing new candidate biomarkers. Among these, we validated *RPL5* LoF mutations as a biomarker of response to selective *MDM2* inhibitors. A dedicated web portal (butterflyvi.unil.ch) enables exploration of all variant annotations and candidate biomarkers. This study highlights the potential and need to expand large-scale functional screenings to empower variant interpretation in the clinic.

## INTRODUCTION

Precision oncology aims at tailoring therapeutic protocols to the features of each tumor and patient. To this purpose, an increasing diversity of molecular assays have entered the clinic. In particular, deep target sequencing of selected gene panels allows to identify cancer-associated variants that may be either clinically actionable through selected inhibitors, or biomarkers of treatment sensitivity. However, each gene and patient can exhibit a wide variety of unique variants, with only a handful of them being functional and/or a clinical biomarker. Variant interpretation consists in determining the functional consequences and clinical implications, if any, of each mutation and it represents today a major bottleneck towards the successful implementation of precision oncology. Currently, variant interpretation largely relies on prior knowledge and expert data curation^1–4^, which are limited and time consuming. On the one hand, predictive approaches have been proposed to prioritize putative functional variants, by leveraging either their recurrence across large patient cohorts^5–7^, evolutionary conservation of the mutated residues^8–10^, or the predicted biochemical and physical alterations to the protein structure, recently adopting cutting-edge deep learning models^11^. While these approaches can rapidly scale to millions of variants, they lack functional validation, which rapidly becomes unfeasible for large numbers of mutations. On the other hand, genome-wide genetic screenings have been used to systematically assess the functional effect of targeting a given gene across multiple in vitro and in vivo models^12,13^, with the largest-to-date examples generated by the Cancer Dependency Map consortium (DepMap)^14–16^. These datasets provide high-throughput functional datasets that could be used by computational models to predict and validate oncogenic dependencies induced by specific variants. Here, we explored the possibility of using results from large-scale functional screenings to systematically classify putative oncogenic and neutral mutations, efficiently providing statistical and functional evidence to interpret thousands of cancer-associated variants.

## RESULTS

The rationale behind the use of genetic screenings for variant interpretation relies on the following hypothesis: if a mutation is functional, i.e. it alters the function of the mutated protein, then selective loss of the corresponding gene has a different effect when the gene is mutated and when it is wild-type (**Fig. 1A**). This hypothesis has been demonstrated for mutations activating known oncogenes. In these cases, oncogenic mutations have been shown to constitute tumor dependencies^17^, which is, tumor cells *depend* on the mutated oncogene and its loss leads to reduced cell proliferation and viability (a.k.a. cell *fitness*), a phenomenon also referred to as oncogene addiction or dependency. Conversely, loss of the same oncogene in cell lines where it does not harbor activating mutations will not affect cell fitness. The identification of oncogene dependencies has provided the rationale for the development of several targeted therapies^18^ and, thus, it has been the main focus of functional genetic screenings^19,20^. However, differential responses to gene loss in mutated vs. wild-type cancer cells can reflect other functional relationships. Beyond oncogene activation, cancer cells rely on the inactivation of tumor suppressor genes, typically through loss-of-function (LoF) variant. As a result, loss of tumor suppressors is expected to increase cell fitness when these genes are wild type, while no fitness changes are expected when they already harbor functional LoF variants. Beyond *oncogene and tumor suppressor dependencies* (OGD and TSD, **Fig. 1B**), two additional types of differential response are possible, which we defined as: *mutation tolerance* (MTO), when loss of the mutated gene increases cell fitness, suggesting the mutation may be disadvantageous to the cell, whereas loss of the wild-type gene has no effect; and, *bypass of essentiality* (BYE), when loss of the wild-type gene decreases cell fitness, which happens for essential genes, whereas loss of the mutated gene has no effect, suggesting that once mutated the gene is no longer essential (**Fig. 1B**). Notably, while oncogene and tumor suppressor dependencies have been previously documented for known cancer genes, the existence and systematic assessment of mutation tolerance and bypass of essentiality among cancer variants is still largely unexplored.

**Fig. 1.**
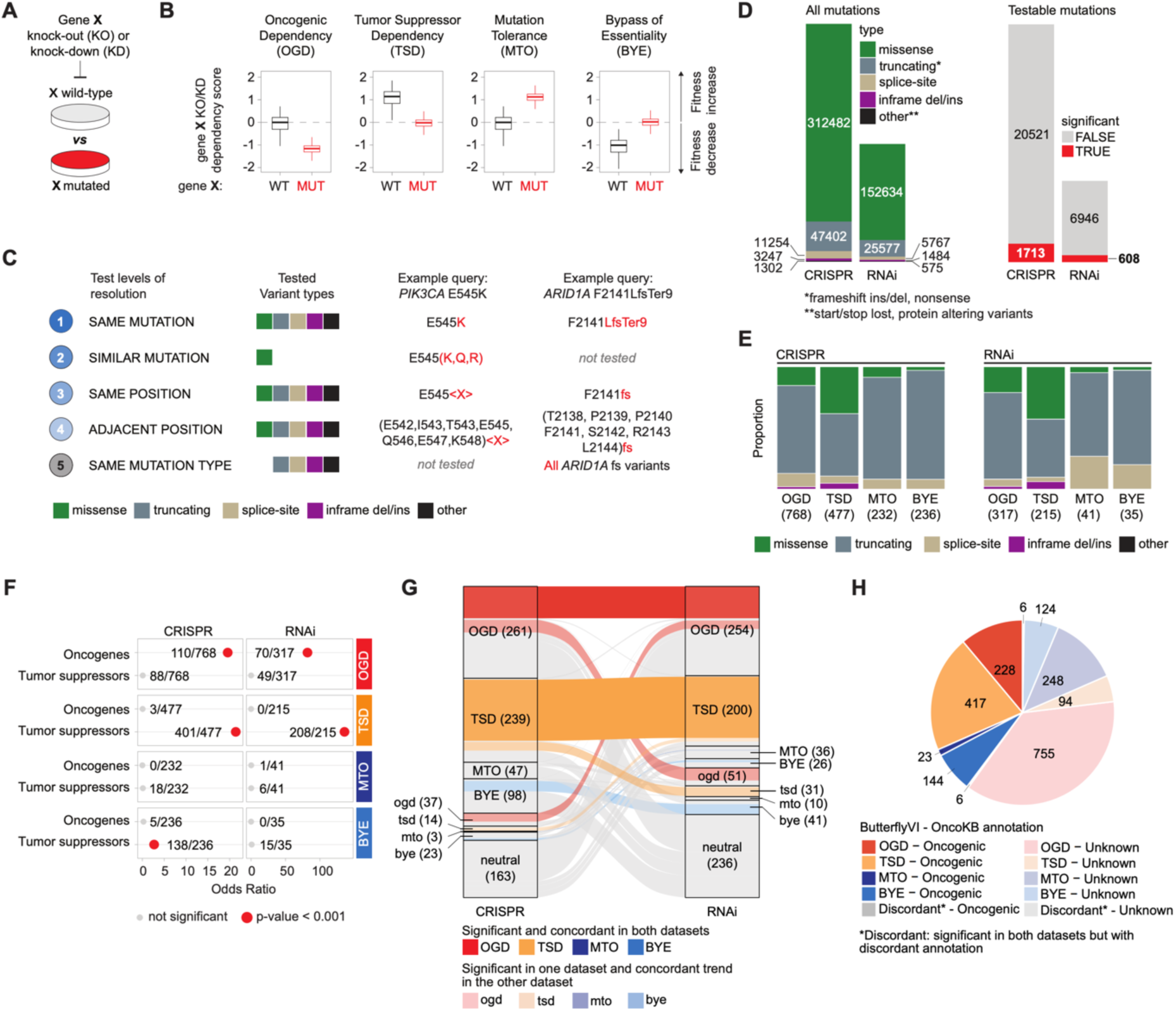
Systematic annotation of mutations in four functional groups. **A**, Schematic of the comparison of the effect of knocking-out/down a given gene in cell lines where that gene is altered (red) and when it is wild type (gray). **B**, Schematics of the four differential responses to gene loss. **C**, Schematics of the five levels of resolution for mutations analysis. **D**, Left: number of mutations per type considered for analysis in CRISPR and RNAi datasets. Right: number of testable and significant mutations in each dataset. **E**, Number of significant mutations in each functional group (OGD, TSD, MTO, BYE) and distribution of mutation types within each group. Colors correspond to the legends in panels C and D. **F**, Enrichment of oncogenes and tumor suppressor genes within each mutation group. The numbers indicate the count of mutations in OG or TSG relative to the total mutations in that group. **G**, Comparison of mutation annotations between the CRISPR and RNAi datasets. The plot includes all mutations that are significant in at least one of the CRISPR or RNAi datasets and are testable in **H**, Distribution of significant mutations based on functional group and OncoKB annotation. The pie chart depicts mutations that are significant in at least one dataset, regardless of testability in the other.

### The ButterflyVI annotation catalog

To unbiasedly test functional dependencies for a large number of unique variants beyond those affecting known oncogenes and tumor suppressors, we integrated and re-annotated molecular data and cell fitness readouts for two large-scale datasets from DepMap: a genome-wide CRISPR knock-out screening dataset^21^, out of which we retained 1178 cell lines, and a genome-wide RNAi knock-down screening dataset^22^, out of which we retained 646 cell lines (**Suppl. Table 1**). In total, these datasets include 375,687 and 186,037 unique variants, respectively. Differential responses to gene loss (i.e., differential dependency scores) between cell lines that harbored a specific mutation for that gene, *altered group*, and cell lines that were *wild-type* were assessed by their Cohen’s D effect size and ANOVA testing, correcting for tumor type. For each test, we required at least 5 cell lines in each group. To test the largest possible number of query variants, we defined 5 levels of resolution, according to which cell lines were considered harboring the query variant if they harbor a mutation that was either exactly the same as the query (**L1** – highest resolution), a “similar” amino-acid substitution, based on a positive BLOSUM62 matrix score^23^ (**L2,** missense mutations only), a mutation occurring at the same residue position (**L3**) or at adjacent positions (**L4**, ± 3 residues), or the same type of mutation of the query variant (**L5** – lowest resolution), with missense mutations being tested only up to L4 (**Fig. 1C** – see Methods). Cell lines were assigned to the wild-type group if they did not harbor any kind of mutations or copy number alterations in the gene of interest. Each variant was tested, and the results retained for the highest resolution where at least 5 cell lines were found in each group. Note that since we tested only mutations that were present in the datasets, at least one cell line harbored each queried variant at L1 resolution. Hence, if a variant was found significant at a resolution lower than L1, we further required that the mean dependency score of the cell lines harboring the query variant at L1 resolution was either (i) closer to the mean dependency score of the altered group than to the mean dependency score of the wild-type group, or (ii) fell below the 0.2 quantile or above the 0.8 quantile of the wild-type dependency score distribution, depending on the direction of the effect (see Methods).

Even allowing for multiple levels of resolution, only 4% and 6% of the total number of mutations were testable, corresponding to 22,234 mutations in the CRISPR dataset and 7,554 mutations in the RNAi dataset, out of which 1,713 and 608, respectively, induced a significant differential response to gene loss (**Fig. 1D**, **Suppl. Table 2**). Once stratified based on the type of differential response, we found that in both datasets most significant variants were either OGD (45% and 52% for CRISPR and RNAi, respectively) or TSD (28% and 35%). Although a minority, in both datasets we identified multiple variants whose differential response to gene loss reflected either mutation tolerance or bypass of essentiality (MTO: 13.5% and 7%, BYE: 14% and 6%) (**Fig. 1E**). In the rest of the study, we will refer to the set of annotations that we derived as the Butterfly catalog for variant interpretation (ButterflyVI), drawing an analogy between the 4 wings of a butterfly and the 4 types of differential response to gene loss.

Consistent with our initial hypothesis, OGD and TSD variants were respectively significantly enriched in oncogenes and tumor suppressors, in both datasets (**Fig. 1F**). Intriguingly, BYE variants were also enriched in tumor suppressors, although only in the CRISPR dataset. Among the 885 variants that were testable in both the CRISPR and RNAi datasets and significant in at least one of the two, 397 (45%) exhibited concordant annotations, and these included annotations across all types of dijerential response, except MTO (**Fig. 1G**). Moreover, of the 2045 variants that were significant in at least one dataset, regardless of whether they were testable in both, 818 (40%) were classified as “oncogenic” or “likely oncogenic” by OncoKB^24^ (**Fig. 1H, Suppl. Fig. 1A-B**), indicating a high agreement between our results and curated experimental and clinical evidence. Importantly, 1221 variants were previously considered of unknown significance and represent novel variant annotations introduced by our study. In a parallel study, OGD were estimated for rare variants using the same datasets but an independent approach (Savino et al. *co-submitted manuscript*). Nicely, OGD estimated by the two studies were significantly concordant, mutually corroborated each other’s results (**Suppl. Fig. 2A-D**).

Most significant dependencies were OGD induced by activating mutations of Ras/Raf oncogenes (*KRAS*, *NRAS*, *HRAS*, *BRAF)* and, to a lesser extent, *PIK3CA*, as well as TSD induced by mutations targeting *TP53*, the significance of which was further driven by the high number of altered cell lines for each variant, and other frequently altered tumor suppressors such as *PTEN* and *RB1* (**Fig. 2A-B**, **Suppl. Fig. 3A-B**). Beyond these well-known oncogenic variants, our analyses revealed significant yet still uncharacterized dependencies, such as OGD site-specific variants at *RHOA* (T19K), *TERT* (H762 splice-site mutation), *IFNA4* (G60E), *ZFP64* (P552/Q553), and *POU3F3* (H311/S312) (**Fig. 2B**).

**Fig. 2.**
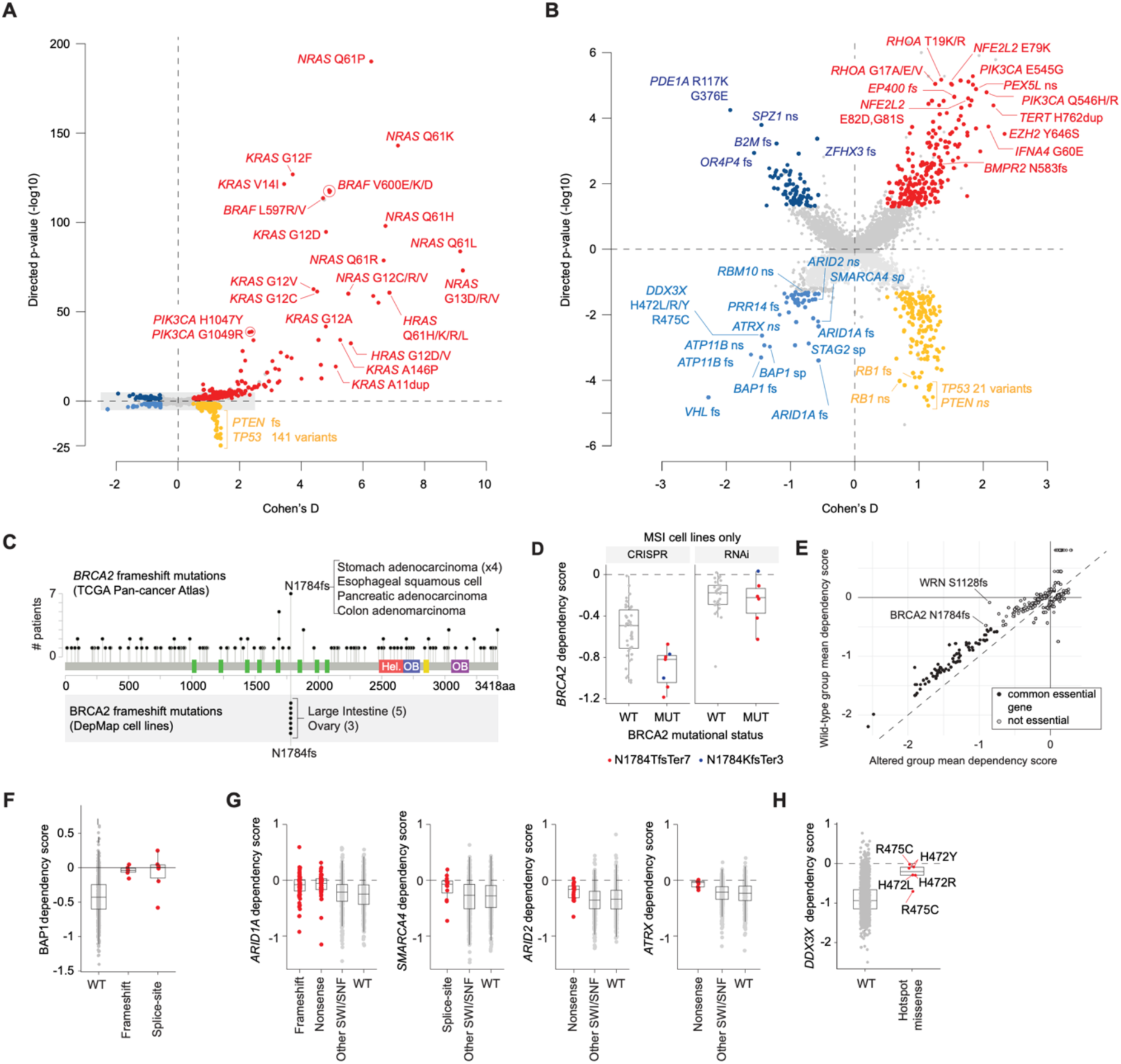
Characterization of the four functional groups of mutations. **A**, Butterfly plot for CRISPR dataset: systematic comparison of gene dependency scores between cell lines carrying a specific mutation and those that are wild type for the corresponding gene. Each point represents a mutation, with its position determined by the Cohen’s D effect size and the associated -log10(p-value) computed at the best testable level of resolution. The p-value is directional: positive for OGD (red) and MTO (dark blue), where wild-type cell lines have mean dependency scores closer to zero than mutant cell lines; and negative for TSD (orange) and BYE (light blue), where mutant cell lines have mean dependency scores closer to zero than wild-type cell lines. **B**, Zoomed-in view of the butterfly plot in panel A (corresponding to the grey-shaded area). **C**, Incidence of BRCA2 frameshift mutations in the TCGA Pan-cancer Atlas (top) and in the DepMap cell lines for the hotspot at position 1784 (bottom). **D,** BRCA2 dependency scores in MSI cell lines that are either wild type for BRCA2 or carry a frameshift mutation at the hotspot position 1784. **E**, Mean dependency score of altered cell lines (x-axis) versus wild-type cell lines (y-axis) in the CRISPR dataset. Each point represents a significant truncating mutation (frameshift or nonsense) classified as OGD, TSD, MTO, or BYE. OGD variants appear in the bottom-left region above the diagonal, indicating increased sensitivity to knockout in altered cell lines compared with wild-type cell lines. **F**, BAP1 dependency scores in the CRISPR dataset for cell lines that are either wild type for BAP1 or carry frameshift or splice-site mutations. **G**, CRISPR dataset dependency scores for four SWI/SNF complex genes in cell lines that are either wild type for the respective gene or carry a BYE mutation. The WT group includes cell lines wild type for all SWI/SNF genes. The “Other SWI/SNF” group includes cell lines that are wild type for the gene considered but carry alterations in other SWI/SNF complex genes. **H**, DDX3X dependency scores in the CRISPR dataset for cell lines that are either wild type for DDX3X or carry a BYE missense mutation in the hotspot region spanning positions 472–475.

Interestingly, a subset of mutations in tumor suppressor genes were also classified as significant OGD events, including *PIK3R1* in-frame and splice region variants and *BRCA2* frameshift hotspot mutation N1784fs. *PIK3R1* variants aligned with two major in-frame mutation clusters that are frequent in uterine endometrial cancer and brain malignancies, as observed in human cohorts from The Cancer Genome Atlas (TCGA) (**Suppl. Fig. 4A**). In the CRISPR dataset, loss of fitness upon knock-out of *PIK3R1* was significant in cell lines harboring in-frame deletions or insertions, and although this signal was more moderate in the RNAi cohort, it remained significant for in-frame insertions (**Suppl. Fig. 4B**). Importantly, these effects were independent of the approach that was used to normalize gene dependency scores in the CRISPR screening (**Suppl. Fig. 4C**). Although these variants are predicted to destabilize inhibitory interactions with *PIK3CA* and favor tumor growth^24^ their functional characterization is still incomplete, and the oncogenic dependency observed here may suggest secondary effects.

*BRCA2* N1784fs mutations were recurrent in gastric tumor samples from TCGA that exhibited microsatellite instability (MSI) (**Fig. 2C**) and, consistently, these variants occurred exclusively in MSI cancer cell lines (**Fig. 2C**). Interestingly, in MSI models, *BRCA2* knock-out was deleterious independently of its mutation status, although dependency scores were significantly lower when the gene harbored the N1784fs hotspot variant (**Fig. 2D**). This difference may be due to gene dosage effects, where loss of one allele through a loss-of-function (LoF) variant increases the dependency of a cell on the remaining allele. A similar case was observed for frameshift mutations targeting the Werner helicase *WRN*, which has been shown to be an MSI dependency^25^. Consistent with a vulnerability induced by loss of heterozygosity, *WRN* frameshift mutations in MSI cancer cells induced a stronger dependency to *WRN* knock-out (**Suppl. Fig. 5**). Deeper investigation of this phenomenon revealed several putative truncating mutations that were classified as OGD, as they induced higher sensitivity to knock-out of genes, whose loss was also harmful in normal cells, but to a lesser degree (**Fig. 2E**). A large fraction of these variants affected common essential genes that increased the gene dependency on the wild-type allele. While deeper investigation will be required for individual cases, these results may reveal novel tumor vulnerabilities, associated with heterozygous LoF variants at common or subtype-specific essential genes^26^.

Putative LoF mutations were also common among BYE variants and affected multiple tumor suppressor genes, seemingly at odds with the observed loss of fitness upon gene knock-out in wild-type cell lines. For example, significant BYE variants comprised multiple LoF mutations affecting the *VHL* and *BAP1* tumor suppressors, which are frequently inactivated in renal carcinoma. Knock-out of *VHL* was deleterious in wild-type cells both at pan-cancer level and within renal cancer cell lines only, consistent with previous experimental evidence demonstrating loss of cell proliferation and/or senescence upon VHL loss^27,28^, while it had no effect in mutated cells, suggesting acquisition of genetic or epigenetic alterations bypassing VHL essentiality (**Suppl. Fig. 6A**). *BAP1* differential responses were observed for both frameshift and splice-region variants (**Fig. 2F**). Among its many functions, *BAP1* is the catalytic component of the polycomb repressive deubiquitinase complex, and it catalyzes deubiquitination of histone 2A (H2A). Interestingly, BYE dependencies were uniquely enriched for LoF mutations affecting histone modifiers, like *BAP1*, and chromatin remodeling factors, in particular, components of the SWI/SNF complex such as *ARID1A*, *ARID2*, *SMARCA4*, and *ATRX* (**Fig. 2B**). Wild-type cell lines or cell lines harboring likely-passenger missense mutations on these genes typically exhibited loss of fitness upon gene knock-out (KO) (**Fig. 2G**, **Suppl. Fig. 6B**), unlike cell lines already harboring LoF events. Similar trends were observed in the CRISPR and RNAi datasets (**Suppl. Fig. 6B-C**) and, importantly, these effects were independent of secondary mutations at SWI/SNF components (**Fig. 2G**, **Suppl. Fig. 6B**), indicating they could not be explained by synthetic lethal interactions^29,30^. Despite the deleterious effect of knocking-out these genes across a wide variety of wild-type cell lines, LoF mutations at these putative tumor suppressor genes are recurrent and considered oncogenic across multiple human tumors^31^. A possible explanation can be found in the “transcriptional numbness” that was recently reported upon KO of epigenetic modifiers in lung cancer and melanoma models^32^, which provided a fitness advantage only in stress conditions. Our results suggest that KO of chromatin modifiers may even be initially deleterious, at least *in vitro*, but eventually promote a cell state where these mutations are tolerated or advantageous. Importantly, we observed such bypass of essentiality across multiple independent models and for multiple genes beyond chromatin remodeling factors.

Indeed, significant BYE variants included recurrent LoF mutations at *ATP11B*, which was among the most significant BYE dependencies, E3 ubiquitin ligases *LTN1* and *UBR2*, and RNA processing genes *RBM10* and *DDX3X*, the latter of which included newly annotated hotspot variants R475C and H472Y/R/L (**Fig. 2H**). As several of these genes have been previously proposed to act as tumor suppressors, our results indicate that their loss may be advantageous only under specific conditions or cell states, which will need to be characterized to understand the oncogenic competence of these variants.

### Biomarker discovery with ButterflyVI annotations

Leveraging our ButterflyVI catalog of putative functional variants, we performed a biomarker discovery analysis to identify predictors of sensitivity or resistance to individual gene loss. Briefly, for each gene, we retained variants annotated as functional in either OncoKB, ButterflyVI, or both, and designed a machine learning strategy based on ElasticNet penalized regression to determine which mutated genes were significantly predictive of the dependency scores in DepMap. Beyond mutated genes, features in the model included: the tumor type of each cell line, microsatellite instability (MSI) status, and gene copy number alterations (see Methods) (**Fig. 3A**). Features were retained as significant predictors if selected among multiple ElasticNet runs (≥50%) and obtaining a mean ElasticNet coefficient greater than .05, in both RNAi and CRISPR datasets (**Fig. 3A**).

**Fig. 3.**
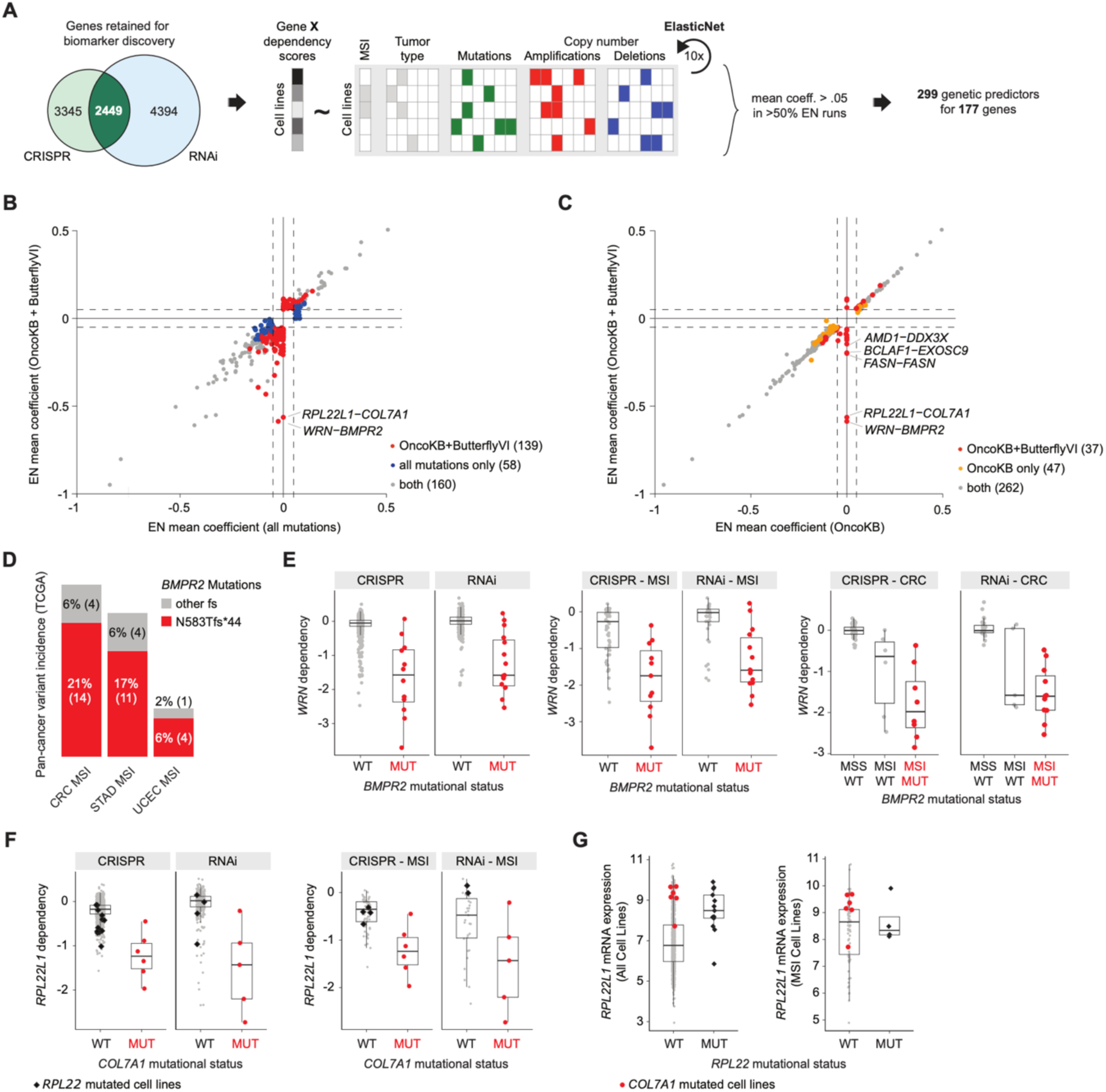
ElasticNet analysis for biomarker discovery integrating functional variants from OncoKB and ButterflyVI. **A**, Schematics of the ElasticNet analysis. **B**, Comparison of ElasticNet scores from the analysis done using all mutations (x-axis) or filtered mutations from OncoKB and ButterflyVI (y-axis). Each point represents a target–biomarker pair. Negative scores indicate sensitivity to the target knockout, while positive scores indicate resistance. **C**, Comparison of ElasticNet scores from the analysis done using filtered mutations only from OncoKB (x-axis) or filtered mutations from OncoKB and ButterflyVI (y-axis). **D**, Frequency of BMPR2 frameshift mutations across tumor types in the TCGA Pan-Cancer Atlas, highlighting the hotspot mutation N583Tfs*44. **E**, WRN dependency scores for cell lines that are either wild type for BMPR2 (or carry a neutral mutation) or carry a functional mutation in BMPR2 (as defined by OncoKB or ButterflyVI). Left: all CLs; centre: MSI CLs only; right: Colorectal Cancer (CRC) CLs only. **F**, RPL22L1 dependency scores for cell lines that are either wild type for COL7A1 (or carry a neutral mutation) or carry a functional mutation in COL7A1 (as defined by OncoKB or ButterflyVI). Left: all CLs; right: MSI CLs only. **G**, Expression levels of RPL22L1 in cell lines that are either wild type for RPL22 or altered (carrying a mutation or deletion).

First, we compared our results with those obtained upon applying the same procedure using only OncoKB annotated variants, or using all variants reported for each gene without any filtering. Even though, ElasticNet coefficients were highly correlated among the different analyses, using all mutations led to a significantly smaller number of genetic predictors (**Fig. 3B**), supporting the relevance of variant annotations and filtering for functional and clinical studies. Interestingly, upon complementing OncoKB annotations with ButterflyVI, we identified new predictors, several of which did not exhibit any association in the OncoKB-only analysis (**Fig. 3C**). Among these, the strongest associations were between mutations at *BMPR2* and *COL7A1* and response to *WRN* and *RPL22L1* knock-out, respectively, both of which have been previously implicated as vulnerabilities in MSI tumors^25^. Notably, the ElasticNet analysis including all mutations did not identify any trend or significant association between these genes (**Fig. 3B**), indicating that this dependency is specifically associated with those variants that were annotated as functional by ButterflyVI. *WRN* encodes for the Werner helicase, which is involved in DNA repair, and it was found as the strongest dependency in MSI tumors, prompting the development of selective inhibitors^33^. *BMPR2* encodes for a bone morphogenetic protein (BMP) receptor that binds TGF-beta ligands and activates downstream SMAD transcriptional regulators. *BMPR2* frameshift mutations were classified as functional by ButterflyVI, including a hotspot frameshift mutation at residue N583, which is recurrent across gastrointestinal MSI human tumors (**Fig. 3D**).

Importantly, differential response to *WRN* knock-out between BMPR2 mutated and wild-type cell lines was independent of MSI status and it was indeed consistently observed across all cell lines, among MSI models only, and among MSI colorectal cancer models, where these mutations were most frequent (**Fig. 3E**). Similar results were obtained for *COL7A1* mutations and *RPL22L1* knock-out. *RPL22L1* encodes a ribosomal large subunit protein that is paralogous to *RPL22,* while *COL7A1* encodes type VII collagen, a structural component of anchoring fibrils in epithelial basement membranes. *RPL22L1* was also reported as a potential MSI dependency, although sensitivity to its loss was attributed to LoF variants in its paralog *RPL22*^25^. Interestingly, cell lines harboring ButterflyVI-annotated *COL7A1* mutations lacked LoF events in *RPL22*, and exhibited a strong dependency to *RPL22L1* knock-out both among all cell lines and within MSI models only (**Fig. 3F**). Consistently, *RPL22L1* was overexpressed in *COL7A1* mutated cell lines, independently of MSI status (**Fig. 3G**). Although residual confounders due to higher order interactions between MSI tumor subtypes and other variants may still exist, our analyses consistently showed increased *WRN* or *RPL22L1* dependency in *BMPR2* or *COL7A1* mutated cell lines, respectively, suggesting that these alterations may confer heightened sensitivity to *WRN* and *RPL22L1* therapeutic inhibition, even among MSI tumors.

Overall, we selected 2449 genes that were testable and exhibited variable response to gene loss in both datasets, and identified 299 genetic features (mutations or copy number alterations) that were significant predictors of response to the knock-out and knock-down of 177 genes (**Fig. 3A**, **Suppl. Fig. 7**, **Suppl. Table 3**). Target genes associated with a high number of predictors included several cell cycle regulators such as *CDK4*, *CDK6*, and *CCND1*, with several predictors corresponding to genes in the same pathway; and oncogenes such as *PIK3CA* and *CTNNB1*, which comprised themselves among the significant sensitivity predictors, consistent with multiple OGD variants at these genes (**Suppl. Fig. 7**). To explore potentially actionable therapeutic biomarkers, we focused on target genes that can be directly or indirectly inhibited through either currently approved cancer therapies or drug compounds in clinical trial. Among these, our ElasticNet model identified multiple known in cis and in trans associations, as well as potentially new biomarkers (**Fig. 4A**). Previously unreported biomarkers included *NOTCH3* amplification that was predictive of sensitivity to *BCL2L1* knock-out (**Suppl. Fig. 8A**), supporting synergistic inhibition of the two oncogenes^34^, *SMAD3* mutations predicting sensitivity to *CDK4* KO (**Suppl. Fig. 8B**), suggesting these variants impair *SMAD3*-mediated cell cycle arrest^35^, and truncating mutations of *RPL5*, which predicted of sensitivity to *MDM2* inhibition (**Fig. 4B**).

**Fig. 4.**
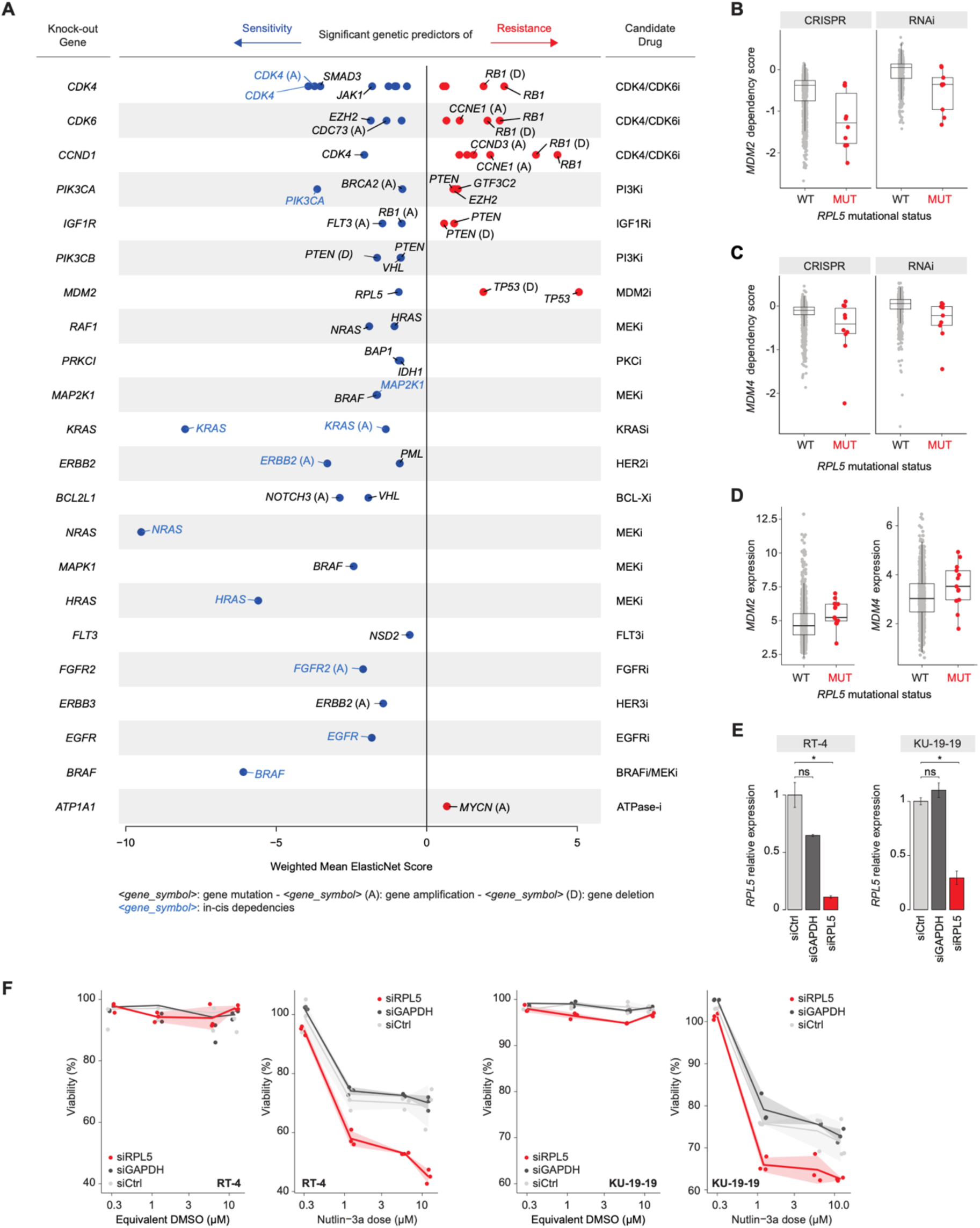
New biomarkers of druggable genes. **A**, Summary plot of ElasticNet biomarkers of druggable genes. Each point shows a biomarker shared between CRISPR and RNAi datasets that was selected at least 5 times out of 10 ElasticNet runs, had a mean coefficient greater in absolute value than .05 and exhibit the same effect direction in both datasets. A negative weighted mean ElasticNet score indicates sensitivity, whereas a positive score indicates resistance. **B–D.** MDM2 (**B**) and MDM4 (**C**) dependency scores, and MDM2 mRNA expression (**D**) in cell lines classified by RPL5 status: wild type (or carrying a neutral mutation) versus carrying a functional mutation (as defined by OncoKB or ButterflyVI). **E**, Relative expression of RPL5 in cells transfected with control siRNA (siCtrl), GAPDH siRNA (siGAPDH), or RPL5 siRNA (siRPL5). Data are presented as the mean ± standard error of the mean (SEM). Differences between groups were evaluated using an unpaired Student’s t-test. The level of statistical significance is indicated by asterisks: ns, not significant (P > 0.05); ** P ≤ 0.01; * P ≤ 0.05. **F**, Cellular viability in response to DMSO (control) or Nutlin-3a treatment at varying doses in RT-4 (left) and KU-19-19 (right) models. Cellular viability is expressed as a percentage. Each point represents a single experimental data point (replicate). The colored lines indicate the mean viability for each knockdown condition (siCtrl, siGAPDH, siRPL5). The shaded bands represent the standard error of the mean (SEM).

*RPL5*, in particular, encodes for the ribosomal protein L5 that has been reported inhibiting MDM2 and activating p53, in response to perturbation of ribosome biogenesis^36^. In the CRISPR dataset, *RPL5* mutations were also predictive of sensitivity to *MDM4* knock-out (**Fig. 4C**) and both *MDM2* and *MDM4* were highly expressed in cell lines exhibiting LoF *RPL5* mutations (**Fig. 4D**). To validate this biomarker, we selected two independent bladder cancer cell lines, RT-4 and KU-19-19, which did not exhibit alterations to the p53 pathway, and we tested their sensitivity to 48 hours treatment with the MDM2 selective inhibitor Nutlin-3a (**Suppl. Fig. 9B**). Next, we inhibited *RPL5* via siRNA in both cell lines and compared response to treatment against scrambled siRNA or an siRNA targeting GAPDH as control (**Fig. 4E, Suppl. Fig. 9C**). In both cell lines, *RPL5* knock-down increased sensitivity to Nutlin-3a (**Fig. 4F**), with differential drug response observed already with low doses of the drug (∼1 μM). Overall, these results indicate that *RPL5* loss sensitizes tumors to MDM2 inhibition and, thus, *RPL5* LoF mutations may represent a novel biomarker of response to MDM2 inhibitors in p53 wild-type tumors.

### The ButterflyVI data portal

To visualize and explore all the results presented in this study we have developed the ButterflyVI Portal (**Error! Hyperlink reference not valid.** The portal is organized in gene-centric pages, which can be searched and accessed by the corresponding HUGO symbol of the gene (**Fig. 5A-B**) and where results are presented in the Variant Interpretation (**Fig. 5C**) and Biomarker Discovery (**Fig. 5D**) pages. Variant Interpretation provides an overview of all tested variants and their corresponding ButterflyVI annotation (**Fig. 5C**), as well as a detailed boxplot representation of the dependency scores in the altered and wild-type group for each variant (**Fig. 5E**). The Biomarker Discovery page summarizes the results of our ElasticNet analysis, reporting for each genes the variants that were found predictive of resistance or sensitivity to loss of that gene (**Fig. 5D**). All data can be downloaded in a table format^a^ and all plots can be saved in common exportable formats (PNG and SVG). ButterflyVI is built using state-of-the-art frameworks. ButterflyVI annotations are stored in a MongoDB NoSQL database for ejicient data access. The front-end application is developed using the Angular framework, and the overview plots are rendered with D3.js. The boxplots are generated using Plotly and ggplot2.

**Fig. 5.**
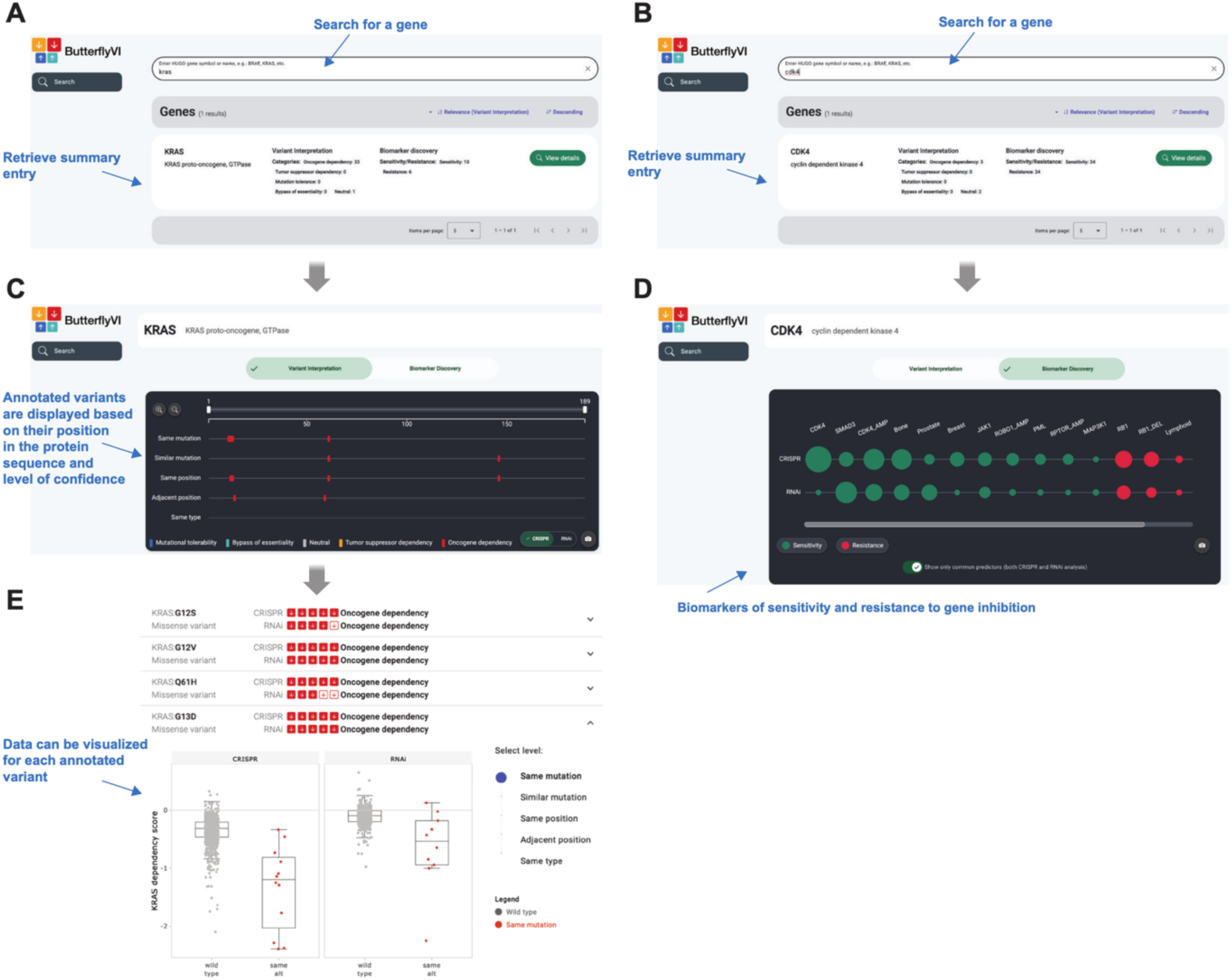
The ButterflyVI data portal. **A-B**, Search page view of the portal. Genes can be queried using their HUGO symbols. Summary gene tabs are displayed underneath for genes with results available from either the variant interpretation or biomarker discovery analyses. **C**, Detailed view of the variant interpretation analysis. Annotated variants are represented as colored rectangles according to their functional category (OGD, TSD, MTO, BYE, or neutral). Their horizontal position corresponds to their location in the protein sequence, while vertical placement reflects the resolution level of testing. A switch allows users to select between CRISPR or RNAi results. **D**, Detailed view of the biomarker discovery analysis. Biomarker of sensitivity (green) or resistance (red) are shown for both CRISPR and RNAi. By default, only common predictors are displayed, but individual dataset predictors can be revealed using the switch at the bottom. Circle size corresponds to the Weighted Mean ElasticNet score. **E**, Overview of all tested variants and their ButterflyVI annotation in CRISPR and RNAi dataset. The number of colored squares indicates the level of resolution (ranging from 5 for same mutation to 1 for the same type level), while the color denotes the functional category of the annotated mutation. Each variant tab can be expanded to show a detailed boxplot of dependency scores in the altered versus wild-type groups. A slider on the right allows adjustment of the resolution level to display the corresponding boxplots.

## DISCUSSION

In this study we presented a systematic framework for functionally annotating cancer-associated variants using two independent genome-wide loss of function screenings. We propose an unbiased classification accounting for four distinct types of differential responses to gene loss, which beyond oncogene and tumor suppressor dependencies, revealed dependencies associated with mutation tolerance and bypass of essentiality. Surprisingly, among variants associated with bypass of essentiality, we found several loss-of-function mutations at tumor suppressor genes, such as *VHL*, *ARID1A*, *ATRX*, and *RBM10*. Although, LoF variants in these genes are recurrent across multiple tumor types, their loss in vitro was deleterious and, for some of them, similar observations were made in vivo. This paradox can be explained by context-specific oncogenic competence where such LoF variants provide an advantage only under certain conditions or cell states^32^, similarly to what has been previously reported for certain oncogenes^37^. Our findings generalized these observations across multiple models and for a broad set of tumor suppressors. We should stress, however, that our analysis solely relies on functional screenings using *in vitro* models, which do not recapitulate the complexity of the tumor microenvironment, both in terms of cell diversity, interactions, and nutrient availability. Moreover, these screenings only measure the impact of gene loss on cell viability, and thus other functionally relevant consequences may not be captured. Despite these limitations, our results and annotations stress the importance of framing functional and clinical studies of variants of unknown significance in the proper context and may allow to discriminate among “universal” tumor suppressors (e.g., *TP53* and *RB1*), which induced tumor suppressor dependencies, and “context-specific” tumor suppressors (e.g., *ARID1A* and *RBM10*), which are associated with bypass of essentiality.

To account for interactions among different types of variants and tumor subtypes, we performed a broad biomarker discovery analysis leveraging OncoKB and ButterflyVI variant annotations. Here, we aimed at identifying genetic variants that predicted sensitivity and resistance to the loss of a broad set of genes, focusing in particular on therapeutically actionable targets. First, this analysis highlighted the importance of filtering variants based on their predicted or validated functional impact. Indeed, biomarker discovery using all variants that were detected for a given gene missed nearly half of significant associations, despite the higher number of retained mutations and therefore greater statistical power. A large fraction of variants within each gene is expected to be neutral and retaining such variants in cancer genomics analyses will inevitably hinder statistical and functional studies. Second, leveraging our annotation catalog we were able to identify novel candidate biomarkers, that could be used to identify patients that could most benefit from a given treatment. Recently, therapeutic inhibition of the Werner helicase WRN has been proposed for microsatellite unstable (MSI) tumors^33^. Here, we showed that tumors exhibiting *BMPR2* loss may exhibit an exquisite sensitivity to this treatment. Similarly, we identified multiple candidate biomarkers for already approved treatments, and, among these, we demonstrated that *RPL5* loss increases sensitivity to Nutlin-3a, a selective MDM2 inhibitor.

Overall, our study demonstrates the relevance of genome-wide functional genomics screenings to enhance variant interpretation and discover therapeutically actionable biomarkers. Nevertheless, these functional assays are still limited in size and, for example, allowed to test only a small fraction of the total number of variants in our cohort. Efforts should be made to scale the generation of these datasets to a larger number and more diverse tumor models, including co-culture assays, tissue explants, and other patient avatar models that could allow to test a broader set of variants and phenotypes. In combination with multi-modal molecular profiling and robust computational models, these approaches will provide an invaluable reference to interpret tumor molecular alterations and translate them into clinically actionable strategies.

## Data availability

The input dataset used to conduct all analyses in this study is available at https://doi.org/10.5281/zenodo.17339253.

## Code availability

The code used to conduct all analyses in this study is available at https://github.com/CSOgroup/ButterflyVI.

## METHODS

### Mutation-specific analysis

#### Datasets

Cell line (CL) data were downloaded from the DepMap portal^15^. Specifically, we used CRISPR-Cas9 gene dependency scores from CRISPRGeneEffect.csv (DepMap 24Q4), RNAi-based dependency scores from D2_combined_gene_dep_scores.csv (DEMETER2 Data v6), somatic mutation data from OmicsSomaticMutations.csv (DepMap 24Q4), and absolute gene-level copy number alterations (CNA) from OmicsAbsoluteCNGene.csv (DepMap 24Q4). Additional metadata, including model annotations and microsatellite instability (MSI) status, were obtained from Model.csv and OmicsSignatures.csv, respectively (DepMap 24Q4). For our analysis, we included only those cell lines that had both gene dependency scores and mutational profiles. This resulted in 1,178 cell lines for the CRISPR dataset and 646 for the RNAi dataset, with 537 cell lines shared between the two. However, CNA data were not available for all selected cell lines: CNA profiles were present for 948 CRISPR-associated and 507 RNAi-associated cell lines. The handling of missing CNA values is described in detail in the subsequent sections.

Mutation data in human cancers, including the TCGA MAF file and corresponding sample annotations, were obtained from Mina *et al.*^38^

#### Sample annotations

We used the *OncotreeLineage* column from the DepMap file Model.csv to define the tumor type of each cell line. Tumor lineages represented by fewer than five CLs in either the CRISPR or RNAi datasets were grouped under the category “other” to ensure statistical robustness.

MSI status was determined using the MSIsensor2 score (*MSIScore* column) from the DepMap file OmicsSignatures.csv, with a threshold of 20% to define MSI-high samples (MSIsensor2 score ≥ 20), in accordance with the recommendations provided by the developers^39,40^.

#### MAF file manual curation

The MAF file from DepMap includes variants annotated with multiple consequence types (e.g., *missense_variant&splice_region_variant*). To enable consistent mutation-specific analysis, we manually curated the data to assign a single variant type to each mutation. In cases of multiple annotations, we applied a prioritization scheme to systematically resolve conflicts, ensuring unambiguous classification for downstream analyses.

Our curation process involved the following steps:

1. Any annotation containing the term “splice” was considered as splice_region_variant.
2. Specific compound annotations involving NMD_transcript_variant were simplified by retaining the primary protein-coding consequence:

- missense_variant&NMD_transcript_variant → missense_variant
- frameshift_variant&NMD_transcript_variant → frameshift_variant
- stop_gained&NMD_transcript_variant → stop_gained
- start_lost&NMD_transcript_variant → start_lost
3. The compound annotation frameshift_variant&start_lost&start_retained_variant was simplified as frameshift variant
4. Any annotation containing the term “start_lost” was considered as start_lost.
5. Any annotation containing the term “start_gained” was considered as start_gained.
6. Any annotation containing the term “stop_lost” was considered as stop_lost.
7. The two remaining compound annotations:

- protein_altering_variant&incomplete_terminal_codon_variant
- coding_sequence_variant&5_prime_UTR_variant were simplified as protein_altering_variant.

#### MAF file OncoKB annotation

We annotated the MAF file using the OncoKB Annotator^41^ to assess the known oncogenic impact of each mutation. Specifically, we performed annotation using the union of the *GenomicChange* and *ProteinChange* options, prioritizing *GenomicChange*, to ensure comprehensive capture of all annotated mutations and to avoid missing variants due to differences in transcript usage or consequence types. In particular, in cases where the *GenomicChange* annotation was labeled as *Unknown*, but the corresponding *ProteinChange* annotation was classified as *Oncogenic*, *Likely Oncogenic*, or *Resistance*, we retained the *ProteinChange* annotation to maximize inclusion of known functionally relevant variants.

#### Selection of mutations to study

To define the set of mutations to analyse we collected all the protein coding region variants present in at least one CL in each dataset. These consist of missense, frameshift, stop gained, splice region, inframe insertion, inframe deletion, start lost, stop lost, and protein altering variants.

#### Copy number alterations

We binarized the copy number profiles of CLs as follows: a gene was considered amplified if the corresponding copy number exceeded 6 copies, and deleted if it was below 1 copy. These thresholds were selected to be consistent with those used in cBioPortal.

#### Test at five levels of resolution

For each selected mutation, we compared the gene dependency scores between CLs that were wild-type for the gene of interest (i.e., without mutations or copy number alterations) and those that carried the mutation. CLs with missing copy number alteration were excluded from the wild-type group. In contrast, inclusion in the mutated group was based solely on mutation status, regardless of copy number information. To evaluate the statistical association between mutational status and dependency score, we performed an ANOVA while controlling for tumor type. This analysis was conducted using the R implementation of ANOVA (function ‘Anova’ from ‘car’ package). For each mutation, we also computed Cohen’s D (D) effect size, which represents the standardized mean difference in dependency scores between wild-type and mutated CLs. A positive D indicates a lower dependency score in the mutated CLs compared to the wild-type, while a negative D indicates a higher dependency score in the mutated CLs relative to the wild-type.

We required a minimum of five mutated CLs to perform the test. To accommodate this, we defined five levels of resolution for the analysis.

1. **L1: same mutation**. This is the highest level of resolution and only CLs with the exact same amino acid substitution were included in the mutated group.
2. **L2: similar mutation** (for missense mutations only). If fewer than five CLs met the previous criterion, we expanded the analysis to include CLs with amino acid substitutions deemed similar based on the BLOSUM62 matrix (i.e., substitutions with a non-negative score).
3. **L3: same position**. If the number of qualifying CLs was still insufficient, or for all non-missense mutations not testable at the same mutation level, we broadened the analysis to include CLs harboring any variant of the same type occurring (for missense) or starting at the same amino acid position of the mutation of interest.
4. **L4: adjacent position**. When needed, we further expanded the analysis to include variants of the same type occurring or starting within ±3 amino acids of the position of interest.
5. **L5: same type** (for non-missense mutations only). Finally, we defined a fifth level of resolution, which included all CLs with the same type of variant anywhere in the gene.

Representative examples of the first three resolution levels are shown in the table below:

| gene | type | same mutation | similar mutation | same position |
| --- | --- | --- | --- | --- |
| PIK3CA | missense_variant | E545K | E545K, E545Q, E545R | Any amino acid substitution at position 545 |
| ARID1A | frameshift_variant | F2141SfsTer59 | - | Any frameshift mutation starting at position 2141 (e.g., F2141LfsTer9) |
| ARID1A | stop_gained | R1989Ter | - | R1989Ter (for stop_gained the same position is equivalent to same mutation) |

Each mutation was tested at the highest possible level of resolution based on data availability.

#### Annotation

We annotated each testable mutation based on its ANOVA p-value and Cohen’s D. In particular, using a cutoff of 0.05 for the p-value and 0.5 for the absolute value of the effect size, we identified five possible scenarios reflecting different effects of the mutation on the cells’ viability. In particular, we annotate each mutation as one of the following:

1. **Oncogenic dependency**: if p<0.05 and D>0.5 and wild-type CLs exhibit a mean dependency score closer to 0 than that of mutated CLs. This case reflects

significant decreased viability of the mutated CLs following gene knock-out/down (whereas wild-type CLs are less affected).

1. **Tumor suppressor dependency**: if p<0.05 and D>0.5 and mutated CLs exhibit a mean dependency score closer to 0 than that of wild-type CLs. This case reflects significant increased viability of the wild-type CLs following gene knock-out/down (whereas mutated CLs are less affected). Note that the direction of the effect size remains the same as in the previous case; what differs is the dependency score’s distribution relative to 0, reflecting increased or decreased cell viability following gene knock-out or knock-down
2. **Mutation tolerance**: if p<0.05 and D<-0.5 and wild-type CLs exhibit a mean dependency score closer to 0 than that of mutated CLs. This case reflects significant increased viability of the mutated CLs following gene knock-out/down (whereas wild-type CLs are less affected).
3. **Bypass of essentiality**: if p<0.05 and D<-0.5 and mutated CLs exhibit a mean dependency score closer to 0 than that of wild-type CLs. This case reflects significant decreased viability of the wild-type CLs following gene knock-out/down (whereas mutated CLs are less affected).
4. **Neutral**: otherwise. In this case no significant difference in the dependency score is observed between the wild-type and mutated CLs.

### Refinement step

To annotate mutations that cannot be tested at the highest resolution L1 (i.e., the *same mutation* level), we introduced a refinement procedure that enhances our confidence at lower resolution levels. If a mutation is deemed significant, the refinement process is initiated. This allows us to distinguish cases where CLs with the same mutation of interest align with the behavior of the broader group of altered CLs, versus cases where the signal is solely driven by CLs with different mutations, while CLs with the same mutation behave similarly to wild-type CLs. Specifically, the refinement process involves two distinct steps.

1. Determine whether the mean dependency score of CLs harboring the same mutation 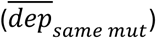 is more similar to that of other altered CLs 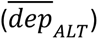 than to wild-type CLs 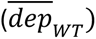. This is assessed by evaluating whether:

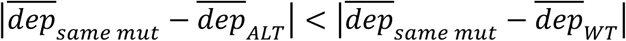

If this condition is met, the annotation is retained; otherwise, the process proceeds to the second step.

2. Determine whether the mean dependency score of CLs with the same mutation lies at the extreme ends of the wild-type CLs dependency score

distribution. Specifically, let 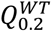 and 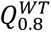 denote the 0.2 and 0.8 quantile, respectively, of the wild type CLs dependency score distribution. We assessed whether:

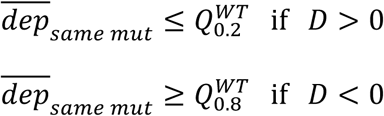

If this condition is satisfied, the annotation is retained; otherwise, the annotation is changed to neutral. This approach allows us to retain mutations for which the mean dependency score of CLs with the same mutation is closer to that of wild-type CLs yet still exhibits a significant effect by lying at the extreme of the wild-type dependency score distribution.

#### Trend of neutral mutations

We focused on all mutations that were significant in either the CRISPR dataset, the RNAi, or both. To better assess the concordance of the annotations between the CRISPR and RNAi datasets, for mutations significant in only one dataset, we assessed their trend in the other dataset using a Cohen’s D threshold of 0.25 and applied the refinement steps. For example, a neutral mutation with a positive Cohen’s D > 0.25 that passed the refinement steps was classified as showing an oncogene or tumor suppressor dependency trend, depending on the relative position of the dependency score distribution compared to zero.

#### Comparison with DAMs from Savino et al

To enable a direct comparison with the results from Savino et al., we first aligned the ProteinChange annotations, as the two analyses were based on differently processed MAF files. For instance, we standardized the representation of stop codons by replacing asterisks (*) with ‘Ter’. Since Savino et al. performed their analysis in a tumor-type-specific manner, a given mutation could appear as significant in multiple tumor types. To ensure consistency in interpretation, we compared their set of unique annotated mutations with our oncogenic dependency results, as their study focused specifically on dependencies with this kind of effect.

#### Selection of the strongest significant annotations

We further filtered the significant mutations to retain only the strongest candidates for subsequent co-alteration analysis. Specifically, we retained mutations that met at least one of the following criteria:

1. Concordant annotation between the CRISPR and RNAi datasets (i.e., the mutation is significant with the same annotation in both datasets, or significant in one dataset and shows the same trend in the other.)
2. Strong effect in at least one dataset, defined as having a p-value < 0.005 and an effect size D > 1.

### Co-alteration analysis

#### Selection of genes to study

We selected the genes that have at least 5 dependent (dependency score < -0.5) and 5 nondependent CLs (dependency score > -0.2) in both the CRISPR and RNAi datasets. This resulted in 2449 genes.

#### Definition of the genomic alteration matrices

We constructed a binary Genomic Alteration Matrix (GAM) capturing the MSI status, tumor type, and the mutational and copy number status of each CL. A gene was considered mutated if it harbored a variant classified as *Oncogenic*, *Likely Oncogenic*, or *Resistance* in OncoKB, or if it had a strong, significant annotation as defined in the previous paragraph. For mutational status, we included all genes with annotations from either OncoKB or TumorScreen. In contrast, for the copy numbers, to limit dimensionality and prevent overfitting, we restricted the set of considered genes. Specifically, we used the union of genes frequently amplified or deleted in cancer (as reported in Sanchez-Vega *et al.*^42^) and genes with mutations annotated in OncoKB. Additionally, for each individual elastic net analysis, we included the respective knocked-out/down gene, if not already included. Amplifications and deletions were encoded as separate binary variables for each gene. We assumed diploid status for CLs with missing copy number data to retain them in the analysis and preserve valuable mutational information. To ensure sufficient statistical power, we included only features altered in at least five CLs. The resulting GAM for the CRISPR dataset includes 1,178 CLs and 669 variables: 1 MSI status, 26 tumor types, 334 mutations, 269 amplifications, and 39 deletions. The GAM for the RNAi dataset includes 646 CLs and 489 variables: 1 MSI status, 21 tumor types, 258 mutations, 186 amplifications, and 23 deletions.

We also defined two alternative GAMs by modifying the criteria for the mutational columns. In the first alternative GAM, a gene was considered mutated based solely on OncoKB annotations (i.e., only if the variant was classified as *Oncogenic*, *Likely Oncogenic*, or *Resistance*) excluding annotations from our method. In the second alternative GAM, a gene was considered mutated if it harbored any somatic mutation, regardless of its annotation. In both cases, we restricted the analysis to the same set of genes included in the original GAM (based on the union of OncoKB and TumorScreen annotations), ensuring that the number of features remained unchanged. The MSI status, tumor type, and copy number alteration columns were kept consistent across all GAMs.

#### Elastic net analysis

For each selected gene, each dataset and each GAM, we performed an Elastic Net (EN) analysis, regressing the continuous gene’s dependency score on the binary variables contained in the GAM. The EN was implemented using the ‘glmnet’ R package, with the mixing parameter set to α = 0.5. The regularization parameter λ was selected via 10-fold cross-validation using the ‘cv.glmnet()’ function, choosing the value that minimized the mean cross-validation error.

To ensure robustness, we repeated the EN analysis 10 times. For each variable, we then computed the mean regression coefficient across the 10 runs, as well as the number of times the variable was selected (i.e., assigned a non-zero coefficient).

#### Filtering of results

To identify the strongest predictors of gene dependency, we selected, for each gene, those predictors with an absolute mean coefficient greater than 0.05 and that were selected in at least 5 out of 10 runs in both datasets and had a concordant sign in CRISPR and RNAi. For each predictor we computed a final score indicating its strength as:

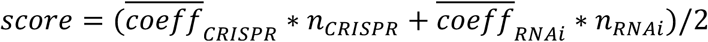

where 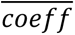 denotes the mean coefficient across the 10 runs, and *n* represents the number of times the variable was selected (i.e., assigned a non-zero value).

#### Experimental validation in bladder cancer cell line models

The human bladder cancer cell lines KU-19-19 and RT-4 (ACC 395 and ACC 412; DSMZ) were cultured in RPMI 1640, GlutaMAX (Gibco, 61870010) supplemented with 10% FBS (Thermo Fisher Scientific, A5256701) and 1% Penicillin-Streptomycin (Thermo Fisher Scientific, 15070063) at 37 °C in a humidified atmosphere with 5% CO₂. FAM-labeled siCTRL No.1 (AM4620), siGAPDH (AM4650), and siRPL5 (ID s56731) (Invitrogen) were reverse-transfected at 25 nM using Lipofectamine RNAiMAX (Invitrogen) in OptiMEM Reduced Serum Medium (Gibco) according to the manufacturer’s instructions. To verify knockdown efficiency, total RNA was extracted using the RNeasy Mini Kit (Qiagen), and reverse-transcribed with the Superscript III First-Strand Synthesis System (Thermo Fisher Scientific). Quantitative PCR (qPCR) was carried out using SYBR Green Real-Time PCR Master Mix (Thermo Fisher Scientific) and gene-specific primers for *GAPDH* (Forward: 5′–CTCTGCTCCTCCTGTTCGAC–3′; Reverse: 5′–ATGGTGTCTGAGCGATGTGG–3′), *RPL5* (Forward: 5′–CCAAATACAGGATGATAGTTCGTG–3′; Reverse: 5′–TTGGCAGTTCGTGTGCATACGC–3′), and the housekeeping gene *HPRT* (Forward: 5′–GTTATGGCGACCCGCAG–3′; Reverse: 5′–ACCCCTTCCAAATCCTCAGC–3′) on a StepOnePlus Real-Time PCR System (Applied Biosystems).

Prior to the main experiment, a Nutlin-3a (Selleckchem, S8059) dose–response assay was performed to determine the sensitivity of KU-19-19 and RT-4 cells, using concentrations based on IC₅₀ values from the Genomics of Drug Sensitivity in Cancer (GDSC) database for KU-19-19 (2.51 µg/mL ≈ 12.18 µM) and RT-4 (2.50 µg/mL ≈ 12.18 µM), including doses below (0.15, 0.3, 0.6, 1.2, 3, 6, and 9 µM) and above (25 and 50 µM) the IC₅₀. For experiments involving *RPL5* knockdown, cells were treated 24 h post-transfection with selected Nutlin-3a concentrations or corresponding DMSO volumes (vehicle control) for 48 h in complete growth medium. At the end of treatment, cell viability was measured using the Cell Counting Kit-8 (WST-8/CCK8 - Abcam, ab228554). Cells were incubated with the ready-to-use reagent for 1 h at 37 °C, and absorbance was measured at 460 nm using a SpectraMax ID3 microplate reader. Absorbance values from blank wells containing only medium were subtracted from sample readings, and cell viability was normalized to untreated controls. Experiments were performed in triplicate for each condition. Data processing and visualization were performed in RStudio.

---

**Suppl. Fig. 1.**
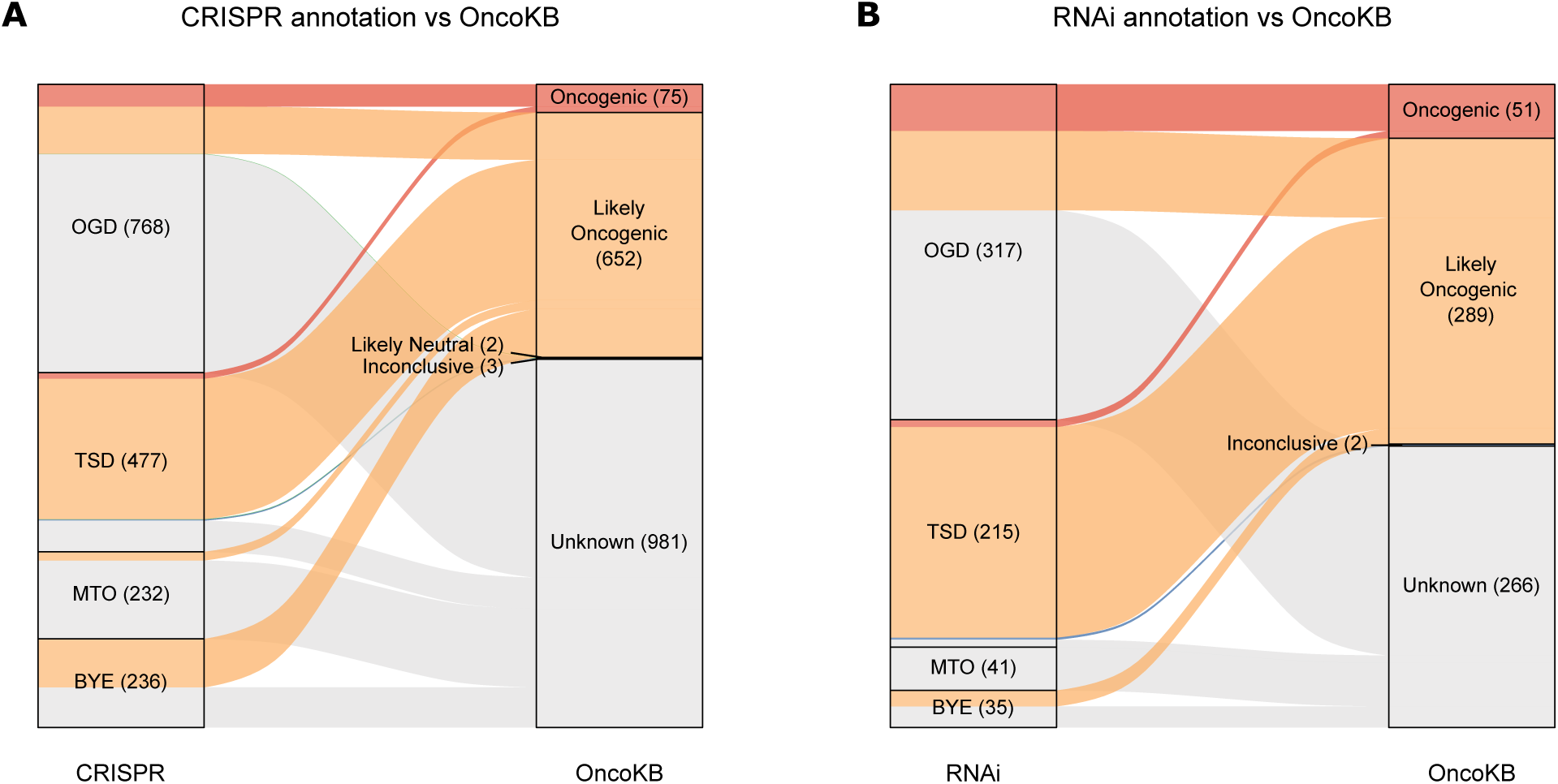
Concordance between single dataset mutation annotation and OncoKB. Comparison of mutation annotations between the CRISPR (**A**) or RNAi (**B**) datasets and OncoKB annotations. The plots include all mutations that are significant in the respective dataset.

**Suppl. Fig. 2.**
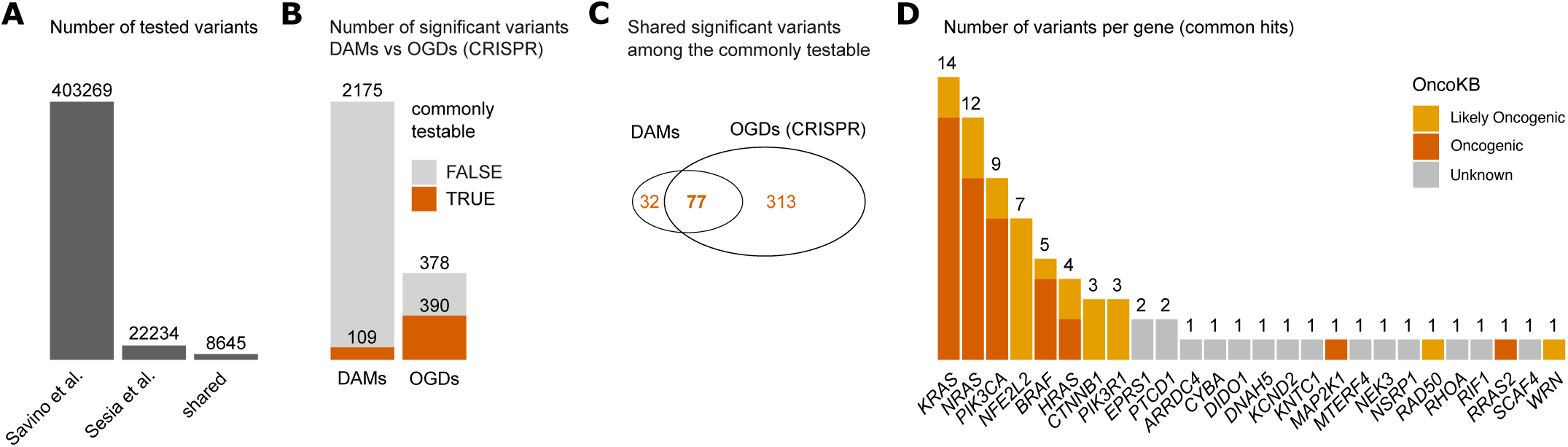
Concordance between OGDs and DAMs from Savino et al. **A**, Number of unique and shared variants tested in the two studies. **B**, Number of significant variants in the two studies. Only OGD variants from our study are considered for the comparison. Variants tested in both studies are shown in orange. **C**, Shared significant variants among the commonly testable. **D**, Number of variants per gene among the shared significant variants. Colors indicate the OncoKB annotation of each variant.

**Suppl. Fig. 3.**
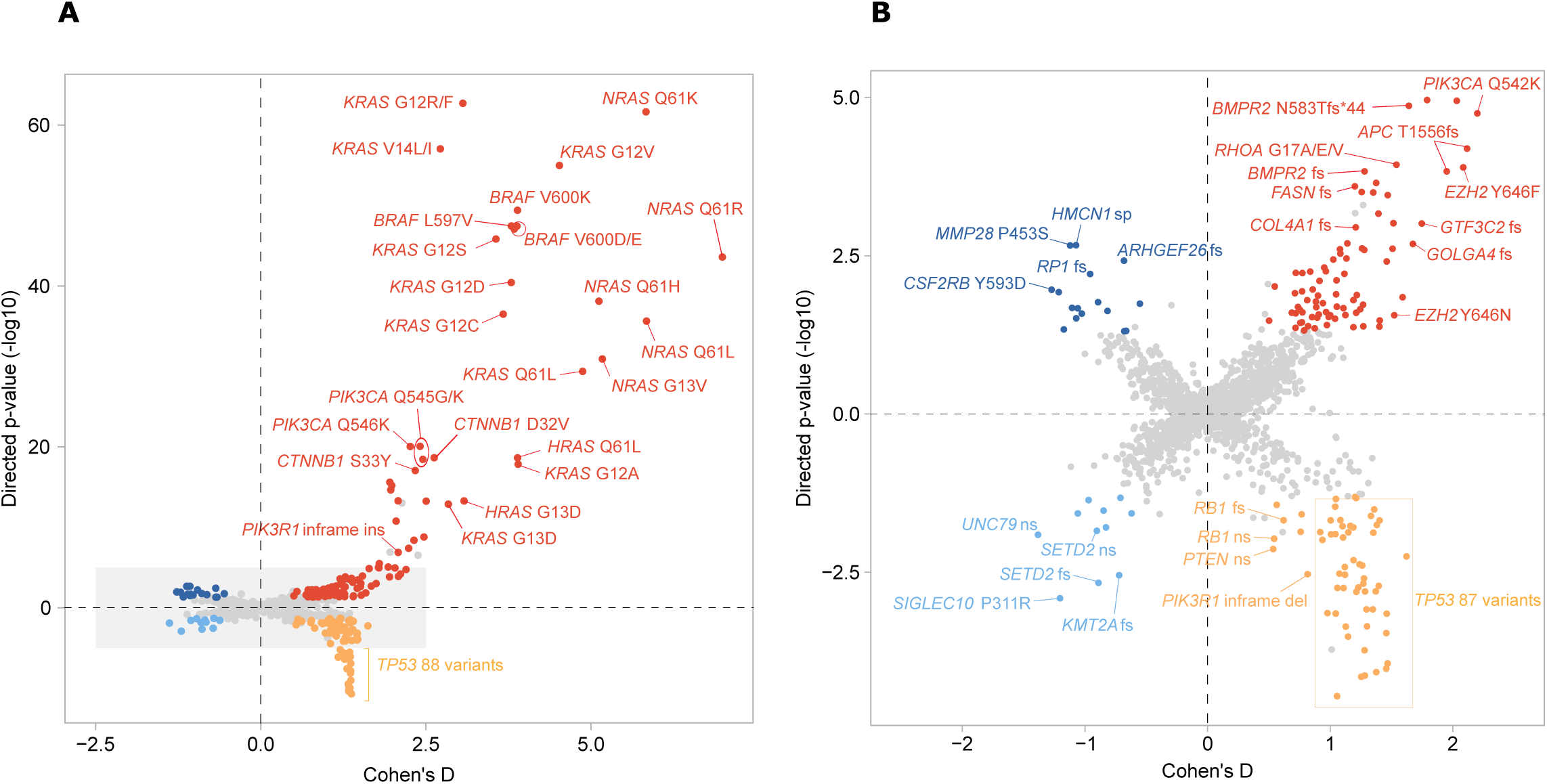
RNAi dataset butterfly plot. **A**, Butterfly plot for RNAi dataset: systematic comparison of gene dependency scores between cell lines carrying a specific mutation and those that are wild type for the corresponding gene. Each point represents a mutation, with its position determined by the Cohen’s D effect size and the associated -log10(p-value) computed at the best testable level of resolution. The p-value is directional: positive for OGD (red) and MTO (dark blue), where wild-type cell lines have mean dependency scores closer to zero than mutant cell lines; and negative for TSD (orange) and BYE (light blue), where mutant cell lines have mean dependency scores closer to zero than wild-type cell lines. **B**, Zoomed-in view of the butterfly plot in panel A (corresponding to the grey-shaded area).

**Suppl. Fig. 4.**
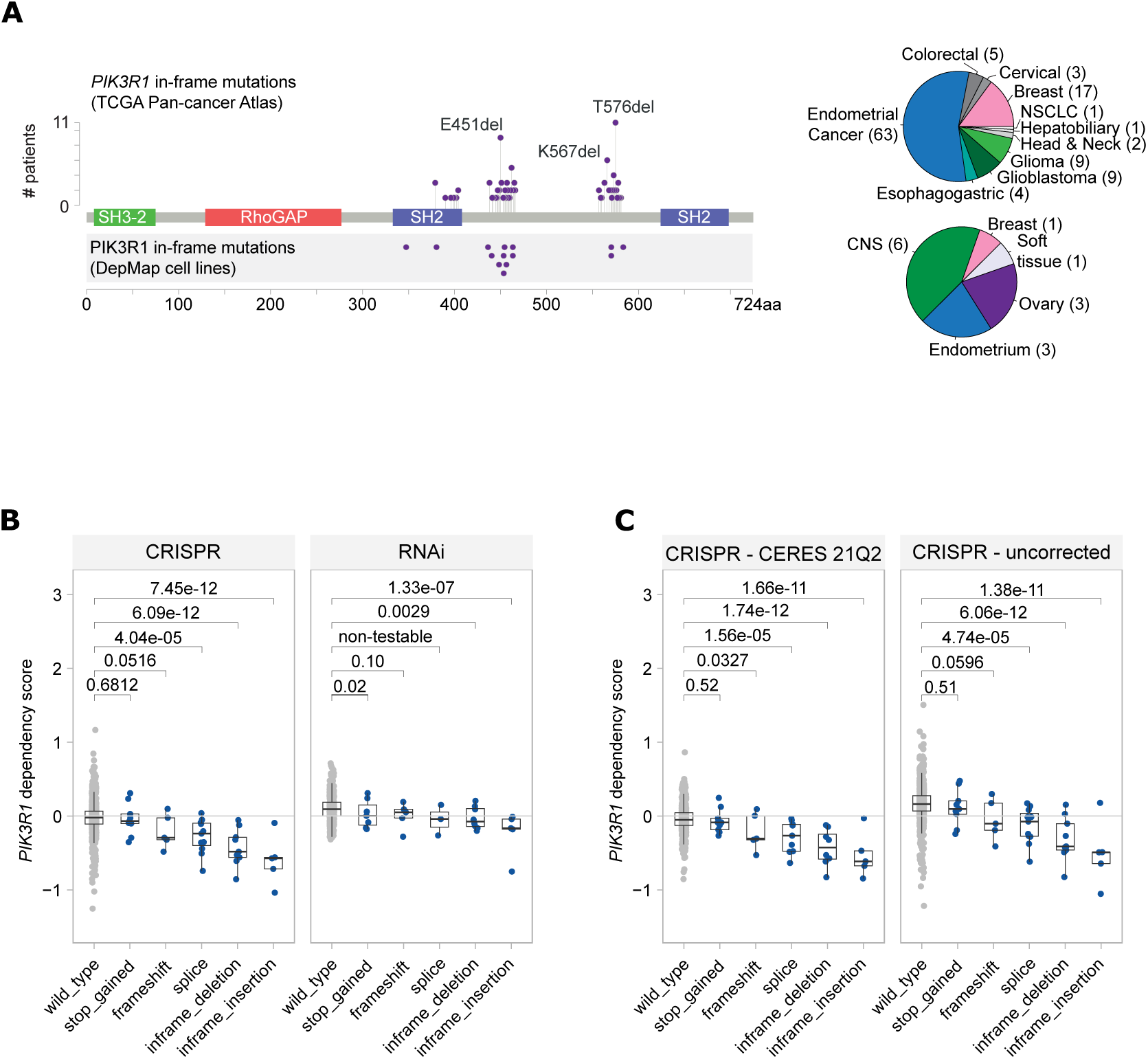
PIK3R1 inframe mutations as OGD. **A**, Left: Incidence of PIK3R1 in-frame mutations in the TCGA Pan-Cancer Atlas (top) and DepMap cell lines (bottom). Right: Tumor type distribution of samples harboring PIK3R1 in-frame mutations. **B**, PIK3R1 dependency scores in cell lines with either wild-type or mutated PIK3R1 for the DepMap data version 24Q4, i.e. the version used for the main analysis. **C**, PIK3R1 CRISPR dependency scores in cell lines with either wild-type or mutated PIK3R1 for the DepMap 21Q2 data version (left) and uncorrected scores from DepMap 24Q4 (right).

**Suppl. Fig. 5.**
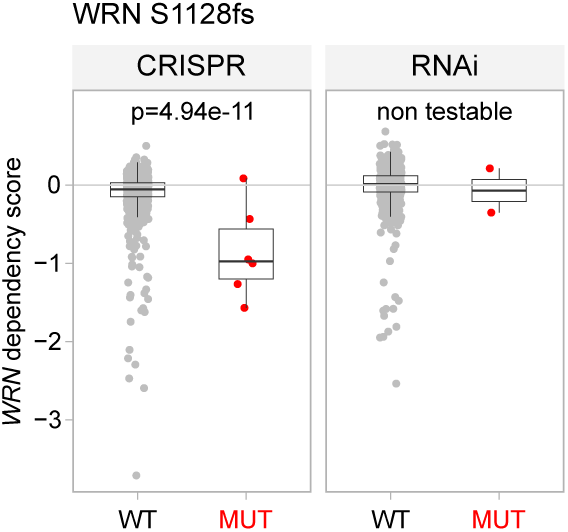
WRN frameshift mutations as OGD. WRN dependency scores for cell lines that are either wild type for WRN or carry frameshift mutations.

**Suppl. Fig. 6.**
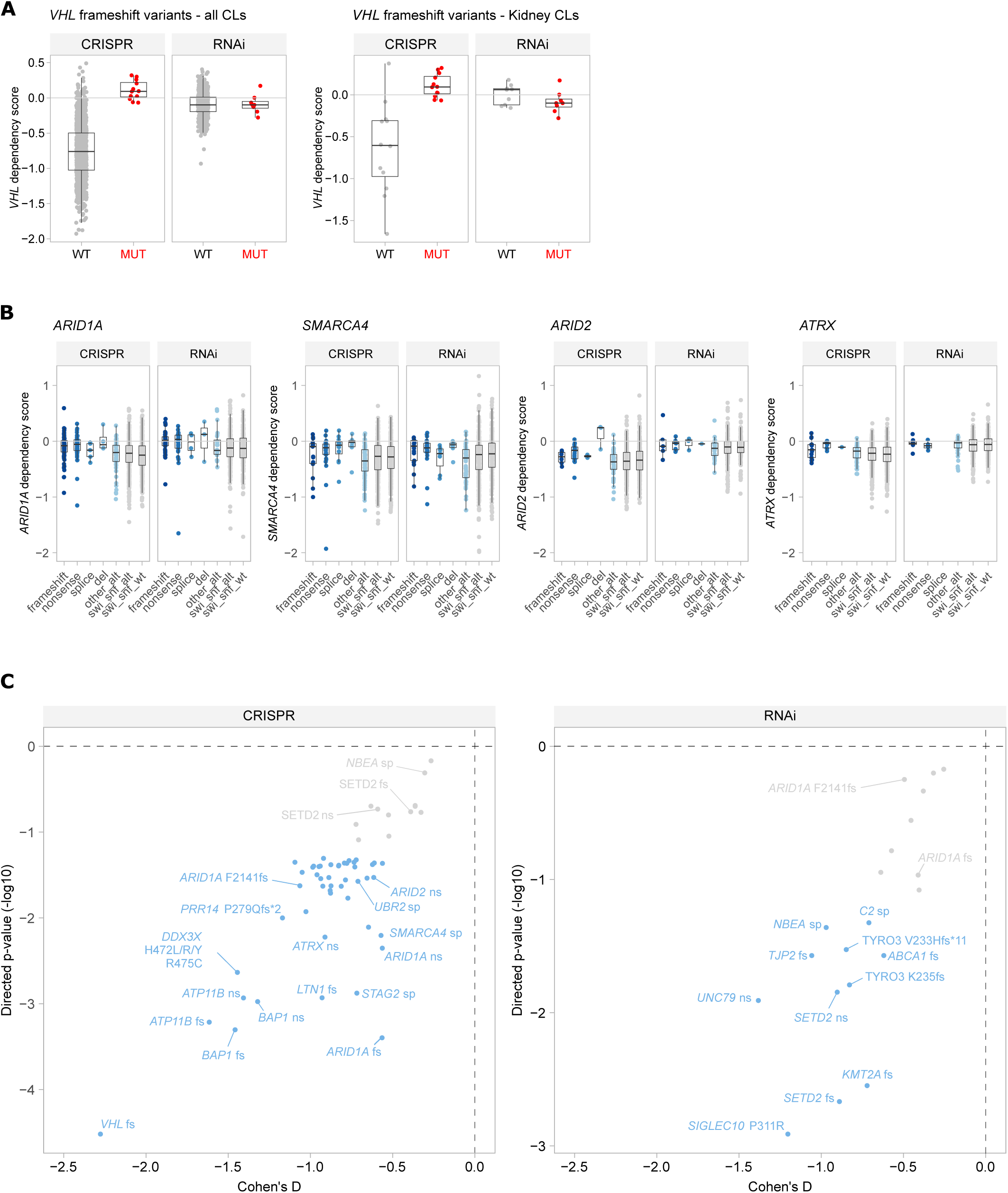
BYE mutations in VHL and SWI/SNF complex genes. **A-B**, VHL dependency scores for all cell lines (**A**) or for kidney cell lines only (**B**) that are either wild type for VHL or carry frameshift mutations. **C**, Dependency scores for four SWI/SNF complex genes in cell lines that are either wild type for the respective gene or carry a specific type of alteration (frameshift, nonsense, splice-site, or deletion). The “swi_snf_wt” group includes cell lines wild type for all SWI/SNF genes. The “swi_snf_alt” group includes cell lines wild type for the gene considered but carrying alterations in other SWI/SNF complex genes. The “other_alt” group includes cell lines carrying other types of alterations (different from the one considered) in the gene of interest. **D**, Zoomed-in view of butterfly plots. BYE mutations are shown in light blue, while neutral mutations with a BYE-like trend (as defined in Methods) are shown in grey.

**Suppl. Fig. 7.**
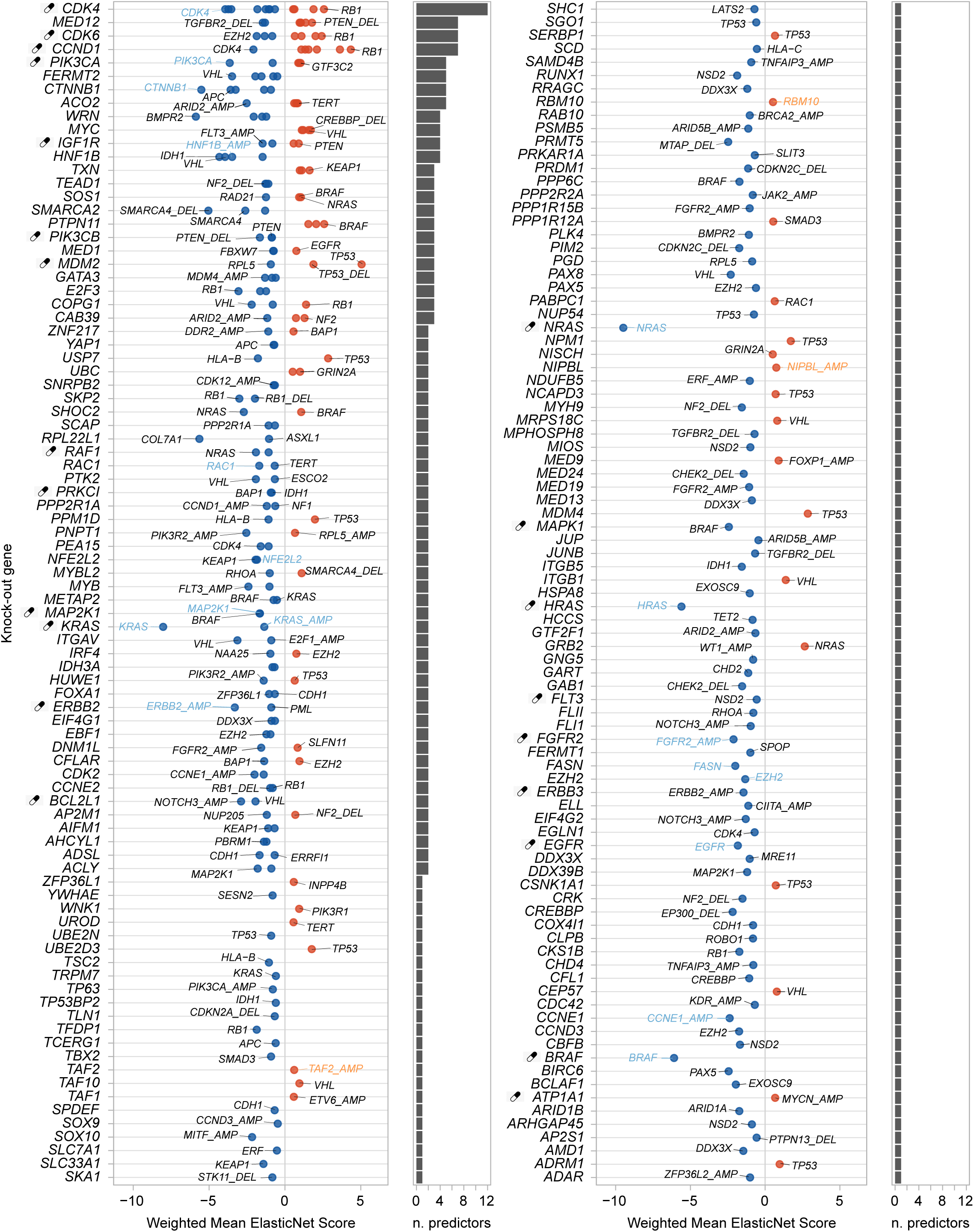
Common biomarkers among CRISPR and RNAi. Summary plot of ElasticNet biomarkers. Each point shows a biomarker shared between CRISPR and RNAi datasets that was selected at least 5 times out of 10 ElasticNet runs, had a mean coefficient greater in absolute value than .05 and exhibit the same effect direction in both datasets. A negative weighted mean ElasticNet score indicates sensitivity, whereas a positive score indicates resistance.

**Suppl. Fig. 8.**
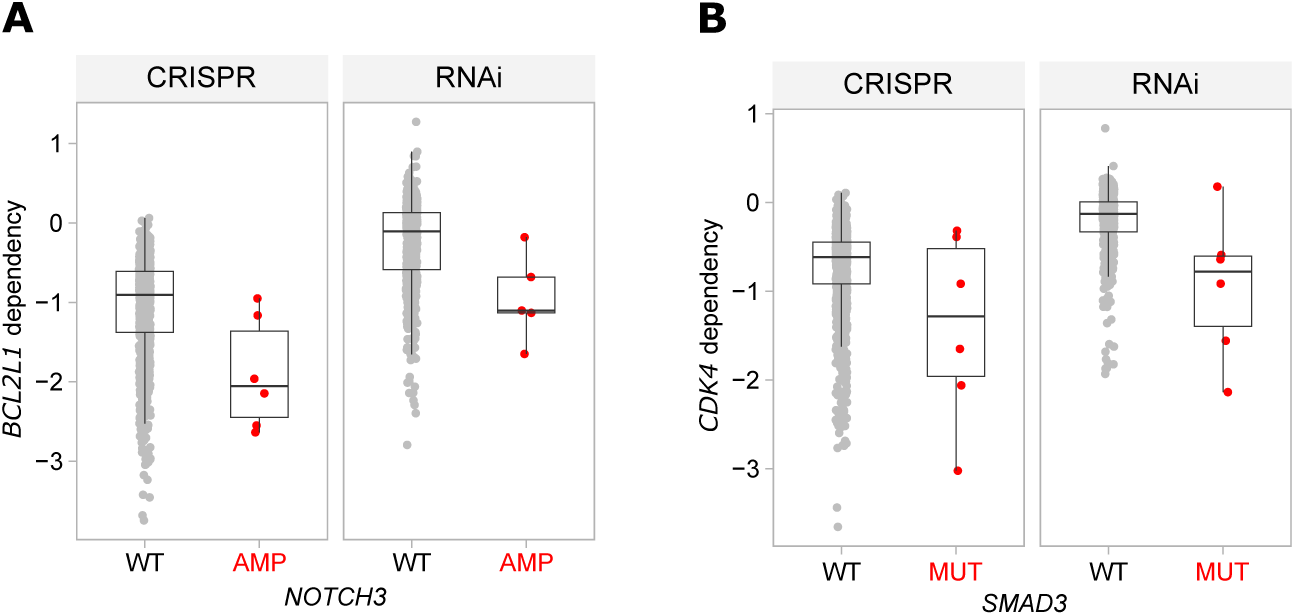
Association between biomarker alterations and druggable gene dependency. **A-B**, Dependency scores for BCL2L1 (**A**) and CDK4 (**B**) in cell lines stratified by biomarker alteration status: wild type (or with a neutral alteration) versus those harboring an amplification (e.g., *NOTCH3* in panel **A**) or a functional mutation (as defined by OncoKB or ButterflyVI; e.g., *SMAD3* in panel **B**).

**Suppl. Fig. 9.**
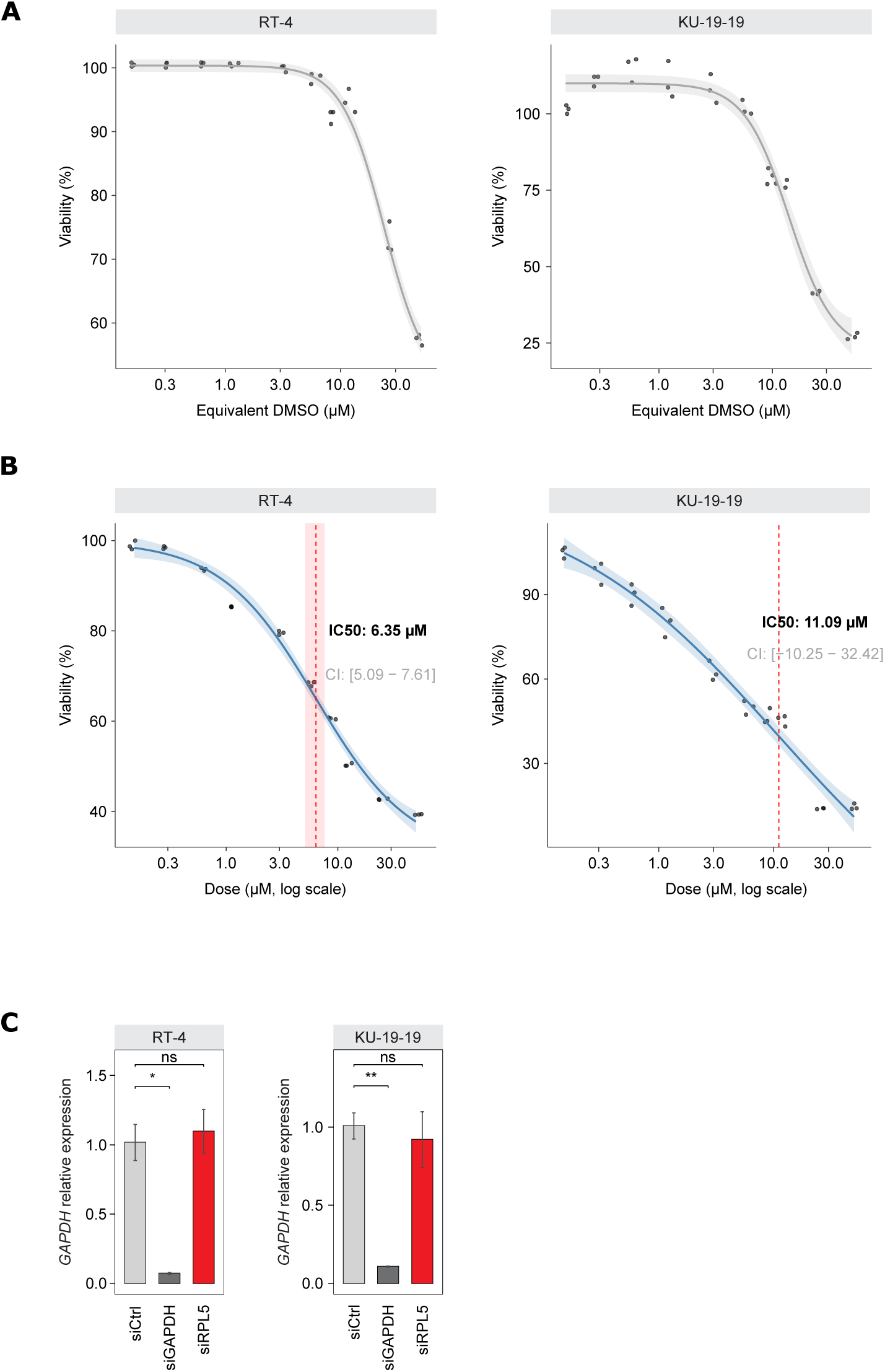
Dose–response curves and siRNA knockdown validation. **A-B**, Dose response curves to control DMSO (**A**) or Nutlin-3a (**B**) treatment on RT-4 and KU-19-19 models. Cellular viability is expressed as a percentage relative to the untreated control. Each data point represents a single experimental data point (replicate). The blue line shows the fit of the four-parameter log-logistic model. The light blue shaded band represents the 95% confidence interval for the model. The vertical red dashed line indicates the calculated IC50, and the pink shaded band represents the 95% confidence interval for the IC50 value. The specific IC50 value and its confidence interval are indicated on the graph. **C**, Relative expression of GAPDH in cells transfected with control siRNA (siCtrl), GAPDH siRNA (siGAPDH), or RPL5 siRNA (siRPL5). Data are presented as the mean ± standard error of the mean (SEM). Differences between groups were evaluated using an unpaired Student’s t-test. The level of statistical significance is indicated by asterisks: ns, not significant (P > 0.05); ** P ≤ 0.01; * P ≤ 0.05.

## Footnotes

a The downloadable files are provided as supplementary material to this manuscript and will be made available on the website upon publication.

